# Development of Triazoles based on AZT and their Anti-Viral Activity Against HIV-1

**DOI:** 10.1101/523225

**Authors:** Daniel Alencar, Juliana Gonçalves, Sofia A. Cerqueira, Helena Soares, Ana Petronilho

## Abstract

We report herein a set of 3’-azido-3’-deoxythymidine (AZT) derivatives based on triazoles and triazolium salts for HIV1 infection. Compounds were tested with □using HIV1 pre-exposure prophylaxis experimental model. All compounds were able to decrease infection and two of them were able to clear almost all the infection, suggesting that these drugs could play an important role in pre-exposure prophylaxis therapies.

## Introduction

The extraordinary development of therapies for Human Immunodeficiency Virus (HIV) treatment over the last decades has extensively reduced the morbidity and mortality associated with the disease. At present, there are over 30 drugs utilized for the treatment of HIV, divided into five main classes, which target different steps in the viral life cycle: 1) viral entry; 2) reverse transcription; 3) integration and 4) viral maturation. However, current therapies are compromised by the rapid emergence of resistant strains, side effects resulting from extended use of drugs and their poor bioavailability^1–3^. The development of more cost effective drugs, with increased efficiency and lower side effects is therefore mandatory. AZT (3’-azido-3’-deoxythymidine) was the first approved drug for the treatment of t HIV^1^. AZT is a nucleoside Reversed transcriptase inhibitor, in which the 3’-OH of the deoxiribose moiety is absent and operates as an obligate chain terminator^1^. Long-term use of AZT is associated with a significant number of side effects, including myopathy, cardiomyopathy, and anemia, among others^2^. AZT is generally used as part of the so-called highly active retroviral therapy (HAART) in combination with lamivudine and abacavir ^4^, still used to date.

Appropriate modification of AZT can be used as a tool to increase the efficiency of the drug and introduce additional functionalities to further increase its rage of activity. The presence of an azide group replacing 3’-OH of the deoxyribose converts AZT in a suitable substrate for further modification by the introduction of triazole groups, known to be active as antivirals in their own right^5,6^. Wang et all showed that modified triazoles can main features which a azole derived AZT must contain to present antiviral activity are a bulky aromatic ring and a 1,5-substitution pattern on the triazole^7^. Nevertheless, the utilization of 1,4 triazoles with sterically demanding substrates also present anti-viral properties, indicating that sterics are probably one of the main driving forces for their activity. In addition, further modifications of AZT within the sugar can also activate or block a specific function, via further modification of the sugar^6^. Herein we report the synthesis of novel AZT derivatives based on triazoles and their antiviral activity against HIV-1, utilizing two methodologies to measure antiviral activity, *e.g.* using infected cells with and without pre-exposure to the drugs.

## Results and Discussion

We synthesized 1,4 triazoles and 1,5 triazoles via 1,3-Huisgen cycloaddition with different substituents and, using both Cu(I) and Ru(II) catalysts, to obtain 1,4 and 1,5 derivatives, respectively^8^.

**Figure 1:**
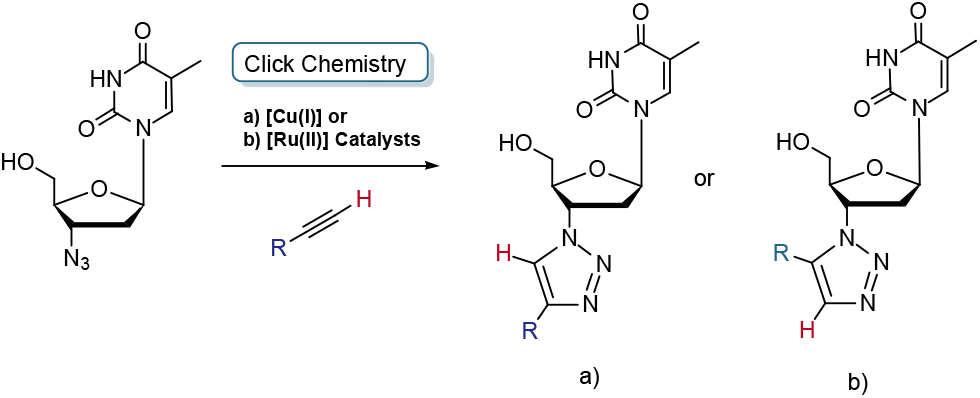
Modification of AZT via Click Chemistry with a) copper(I) and b) ruthenium (II) Catalysts

For 1,4-triazoles, AZT reacts with the corresponding alkyne with the Cu(I) catalyst (5%) at 100 °C for 2h, in a mixture of tert-butanol:water (1:1). For 1,5-triazoles require somewhat longer reaction times (24 hours) using a similar protocol with Ru(II)(C_5_Me_5_)(PPh_3_)Cl (5%) at 100 °C in dioxane. In both cases, purification via silica gel chromatography is required. Compounds are obtained in good to moderate isolated yields (50 to 80%).

**Scheme 2:**
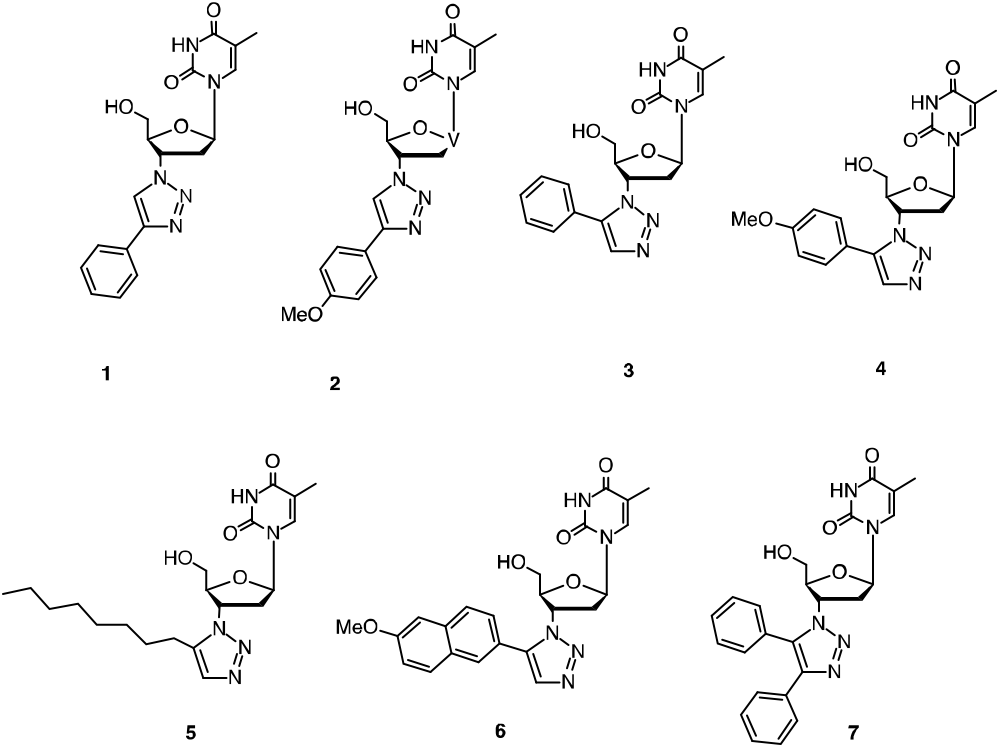
Representative 1,4- and 1,5 triazoles synthesized in this work, compounds 1-7.

Aiming to understand how the anti-viral activity is linked to the two methods utilized in this work, we synthesized two standard 1,4- and 1,5-phenyl derivatives, **1** and **3**, as well compound **6**, previously reported, for comparative purposes^7^.

To confirm the structures of the compounds, NMR studies (^1^H and ^13^C, 1D and 2D) were conducted. The formation of the 1,4 and 1,5 substituted triazole is clearly indicated with the presence of a singlet, corresponding to the newly formed CH bond. For 1,4 triazoles, this singlet is located between 8.5-9.0 ppm, while for 1.5 triazoles, it resonates at somewhat higher field, between 7.5-8.0 ppm. For compound **7** ^13^C clearly indicates the formation of the triazole ring with the presence of two quaternary carbons at 125.2 and 127.3 ppm.

Further functionalization of the AZT triazole derivative via quaternization of N3 of the triazole ring^9^ is possible. This synthetic variability could in principle improve physiological solubility, due to the existence of a delocalized positive charge at the triazole and formation of a salt. Thus, we synthesized the corresponding triazolium salts derived from compounds **1-7** using methyl iodide or methyl triflate, to obtain the triazolium salts **8** and **9**. The compounds were obtained in moderate yields (50-60%). Their formation was also confirmed by NMR studies and MS. In the ^1^H NMR, and when compared to the corresponding triazoles, a significant downfield shift of *circa* 1 ppm is also observed for the aromatic C-H of the triazole ring. This feature is diagnostic of N3 quaternization, and formation positive charge on the nitrogen that is delocalized within the triazole ring, inducing the deshielding of the H_trz_ proton. In addition, a new singlet appears in the region δ 3.5-4.5 ppm indicative of the presence of the methyl group at N3.

**Scheme 3:**
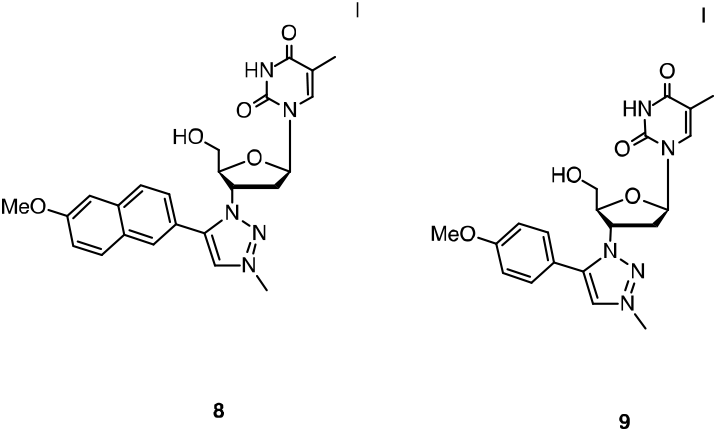
triazolium salts **8** and **9** synthesized by methylation of compounds **4** and **6**.

## Anti-Viral Activity

Next, we determined the ability of the compounds synthesized to block HIV-1 replication in human primary CD4 T cells isolated from peripheral blood and finally of healthy donors ^10^.

Two main methods were tested as depicted in Fig. 4, in the first one we tested their anti-retroviral capacity once HIV-1 replication had already initiated (method 1), while in the second methodology we tested their ability to act as pre-exposure prophylaxis treatment we pre-treated CD4 T cells with AZT derivatives for the 18 hours prior to HIV-1 infection (method 2).

**Figure 4:**
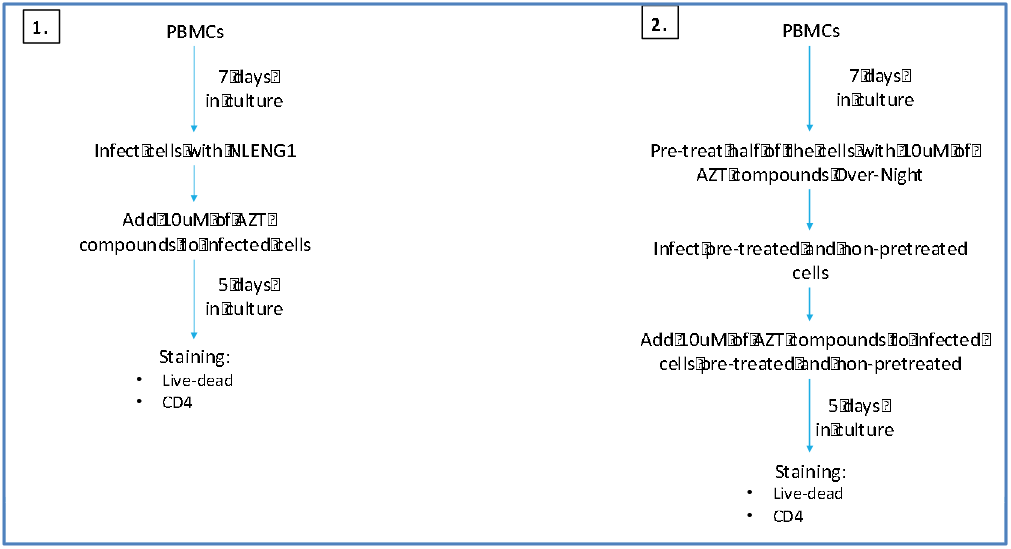
Schematic representation of methodological approach. Human peripheral blood cells were isolated from whole blood through gradient centrifugation^11^. Primary CD4+ T cells were infected with a GFP-tagged NLENG1-IRES virus for 5 days^12^ AZT-derivatives were either added post-(methods 1) or 18 hours pre-infection. At the end of the 5 days HIV-1 replication was assessed by comparing the frequency of HIV-1 infected cells in non-treated versus AZT-derivatives treated conditions by flow cytometry.

The percentage of HIV-1 infected cells were determined in live CD4 T cells, according to the gating strategy outlined in Fig. 5. Part of the compounds was tested with these two methodologies, and the results can be found bellow.

**Figure 5:**
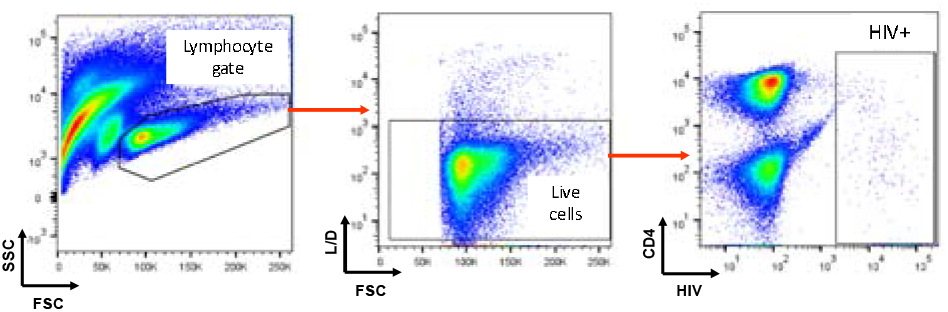
Flow cytometry gating strategy. We incubated HIV-1 infected primary CD4+ T cells with the live/dead fixable Viability Dye eFluor 780 (eBioscience). The frequency of HIV-1 infected primary CD4^+^ T cells was determined by gating on GFP^+^ live primary CD4^+^ T cells.

**Figure 6:**
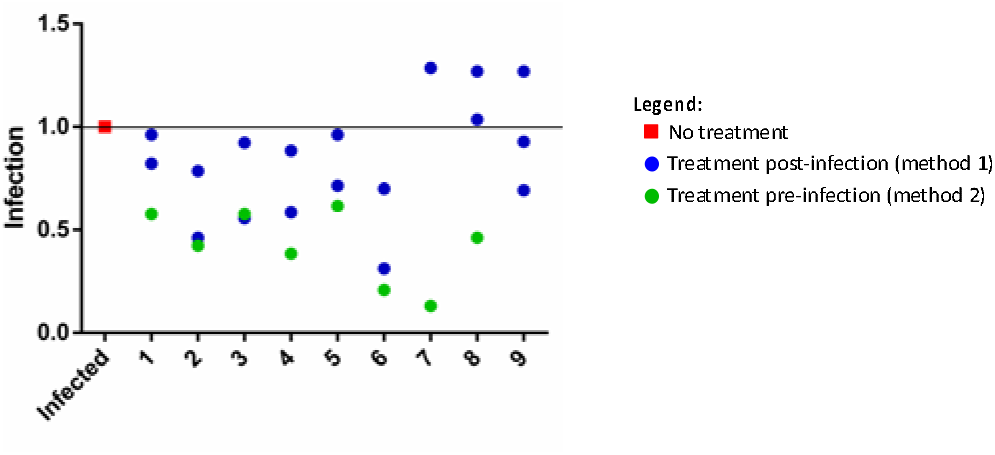
AZT-derivatives exhibit pre-exposure prophylaxis potential. Plotted fold-change in the frequency of HIV-1 infected CD4^+^ T cells upon treatment with different AZT-derivatives.

We found that AZT derivatives have modest effects in HIV-1 replication once primary round of infection had been established (method 1). Nevertheless, compounds **2** and **6** seem to be able to partially halt ongoing HIV-1 ongoing replication. We obtained far more promising results with AZT derivatives in our pre-exposure prophylaxis experimental model (method 2). In this latter experimental approach, all compounds tested were able to decrease HIV-1 infection rates for at least one half, and remarkably compounds **6** and **7** were able to clear almost all infection.

Regarding the activity of 1,4-versus 1,5-triazoles, these results indicate that there is no significant difference between their antiviral activity for method 1, with the two most active compounds are compounds **2** and **6**. As for the activity of the triazolium salts, our observations indicate that there methylation decreases antiviral activity. These preliminary results are in line with previous reports regarding the activity of 1,5-triazoles derived from AZT, with compound **6** reported by Wang^7^ presenting high antiviral activity for the two methods utilized.

## Conclusions

Despite the efficacy of AZT, the search of other HIV-1 reverse transcription inhibitors has continued due to AZT poor bioavailability at the sites of HIV-1 replication (lymphoid organs), the emergence of HIV-1 resistant strains and secondary effects. Here we describe novel AZT derivatives modified with triazoles that are particularly efficient if provided prior to HIV-1 infection. All compounds were able to decrease infection and two of them, compounds **6** and **7**, were able to clear almost all the infection. Our preliminary results suggest that these drugs could play an important role not only in HIV-1 treatment but also in pre-exposure prophylaxis therapies. Further derivatization of AZT is currently being pursued and will be reported in due course.

## Acknowledgements

We thank Cláudia Andrade for Flow Cytometry support, the National Institutes of Health AIDS Reagent Program for providing the human IL-2. The NLENG1-IRES GFP-tagged HIV-1 virus was generously provided by David N. Levy.

## EXPERIMENTAL PROCEDURES

### Compound 1

A suspension of 3′-azido-3′-deoxythymidine (800 mg, 3.00 mmol), phenylacetylene (395 μL, 3.60 mmol), CuSO_4_ (29 mg, 0.18 mmol) and sodium ascorbate (357 mg, 1.80 mmol) in H_2_O/tert-butanol (30 mL 1:1 v/v) was stirred at 100 °C for 2 hours. After cooling, the solid residue was filtered off and washed with Et_2_O. The product was then purified by silica gel chromatography (CHCl_3_: MeOH 10:1 v/v). Compound **1** was obtained as a white powder (922 mg, 83%). ^1^H NMR (400 MHz, DMSO-*d*6) δ 11.38 (1H, s, NH), 8.79 (1H, s, H_trz_), 7.90 − 7.81 (3H, m, H6, H_Ar_ and H_Ar_), 7.47 (2H, t, ^3^*J*_HH_ = 7.6 Hz, H_Ar_ and H_Ar_), 7.36 (1H, t, ^3^*J*_HH_ = 7.6 Hz, H_Ar_), 6.46 (1H, dd, ^3^*J*_HH_ = 6.6 Hz, H_1′_), 5.41 (1H, dt, ^3^*J*_HH_ = 8.8, 5.2 Hz, H_3′_), 5.31 (1H, dd, ^3^*J*_HH_ = 5.2 Hz, OH), 4.29 (1H, dt, ^3^*J*_HH_ = 5.2, 3.6 Hz, H_4′_) 3.77 − 3.65 (2H, m, H_5′_), 2.85 − 2.67 (2H, m, H_2′_), 1.83 (3H, s, CH_3_). ^13^C{^1^H} NMR (100 MHz, DMSO-*d*6) δ 164.2 (C=O), 150.9 (C=O), 147.0 (C_q_), 136.7 (C6), 131.0 (s, Cq_Ar_), 129.4 (2CH_Ar_), 128.4 (CH_Ar_), 125.6 (2CH_Ar_), 121.4 (C_trz_), 110.1 (C5), 84.8 (C4′), 84.3 (C1′), 61.2 (C5′), 59.8 (C3′), 37.6 (C2′), 12.7 (CH_3_).

### Compound 2

3′-azido-3′-deoxythymidine (100 mg, 0.37 mmol), phenylacetylene (80 μl, 0.75 mmol), and Cp*RuCl(PPh_3_)_2_ (0.05 eq.) were dissolved in dioxane (2 ml) under N2 atmosphere, the resulting mixture stirred at 80°C for 48 hours. After cooling the solvent were then removed under reduced pressure and the solid residue was purified by silica gel chromatography (DCM:MeOH 10:1 v/v). Compound **5** was obtained as a brown powder (116 mg, 85.5%). ^1^H NMR (400 MHz, DMSO-d6) δ 11.36 (1H, s, NH), 7.92 (1H, s, H_trz_), 7.78 (1H, s, H_Ar_), 7.60-7.53, (5H, m, H_Ar_), 6.57 (1H, dd, ^3^*J*_HH_ = 6.8 Hz, H1′), 5.25 (1H, dd, ^3^*J*_HH_ = 5.2 Hz, OH), 5.19-5.15 (1H, m, H3′), 4.40 (1H, dt, ^3^*J*_HH_ = 3.6, 3.2 Hz, H4′), 3.63-3.47 (2H, m, H5′), 2.66-2.57 (2H, m, H2′), 1.78 (3H, s, CH_3_). ^13^C{^1^H}NMR (100 MHz, DMSO-*d*6) δ 164.1 (s, C=O), 150.9(C=O), 138.3 (Cq_Ar_), 136.5 (CH_Ar_), 133.3 (CH_trz_), 129.6 (5C, CH_Ar_), 126.8 (CH_Ar_), 110.1 (C_Ar_), 85.3 (C4′), 85.1 (C1′), 61.8 (C5′), 58.1 (C3′) 38.1 (C2′), 12.7 (CH_3_).

### Compound 3

3′-azido-3′-deoxythymidine (100 mg, 0.37 mmol), ethynylanisole (97 mg, 0.75 mmol), and Cp*RuCl(PPh_3_)**2** (0.05 eq.) were dissolved in dioxane (5 ml) under N2 atmosphere, the resulting mixture stirred at 80°C for 48 hours. After cooling the solvent was removed under reduced pressure and the solid residue was purified by chromatography (DCM:MeOH 10:1 v/v). Compound **3** was obtained as a brown powder (47 mg, 31.3%). ^1^H NMR (400 MHz, DMSO-d6) δ 11.36 (1H, s, NH), 7.85 (1H, s, H_trz_), 7.79 (1H, d, ^4^*J*_HH_ = 0.8 Hz, H_Ar_), 7.47 (2H, d, ^3^*J*_HH_ = 8.0 Hz, H_Ar_), 7.12 (2H, d, ^3^*J*_HH_ = 8.0 Hz, H_Ar_), 6.57 (1H, dd, ^3^*J*_HH_ = 6.4 Hz, H1′), 5.25 (1H, dd, ^3^*J*_HH_ = 4.8 Hz, OH), 5.16-5.14 (1H, m, H3′), 4.38 (1H, dt, ^3^*J*_HH_ = 3.6, 3.2 Hz, H4′), 3.83 (3H, s, O-CH_3_), 3.63-3.49 (2H, m, H5′), 2.62-2.51 (2H, m, H2′), 1.79 (3H, s, CH_3_). ^13^C{^1^H}NMR (100 MHz, DMSO-*d*6) δ 164.1 (C=O), 160.6 (C_Ar_), 150.9 (C=O), 138.1 (C_q_), 136.5 (CH_Ar_), 133.1 (CH_Ar_), 131.1 (2CH_Ar_), 118.7 (CH_Ar_), 115.0 (2CH_Ar_), 110.1 (Cq_Ar_), 85.3 (C4′), 85.1 (C1′), 61.8 (C5′), 58.6 (C3′), 55.7 (O-CH_3_), 38.2 (C2′), 12.7 (CH_3_).

### Compound 4

A suspension of 3′-azido-3′-deoxythymidine (100 mg, 0.375 mmol), ethynylanisole (60 mg, 0.450 mmol), CuSO_4_ (3.5 mg, 0.022 mmol) and sodium ascorbate (45 mg, 0.225 mmol) in tert-butanol (5 mL) was stirred at 100 °C for 2 hours. After cooling the solid residue was filtered off and washed with Et_2_O. The solid residue was purified by silica gel chromatography (DCM:MeOH 10:1 v/v). Compound **4** was obtained as a white powder (50 mg, 33.3%).

^1^H NMR (400 MHz, DMSO-d6) δ 11.37 (1H, s, NH), 8.67 (1H, s, H_trz_), 7.85 (1H, d, ^4^*J*_HH_ = 0.8 H_Ar_), 7.785 (2H, d, ^3^*J*_HH_ = 8.8 Hz, H_Ar_), 7.045 (2H, d, ^3^*J*_HH_ = 8.8 Hz, H_Ar_), 6.45 (1H, dd, ^3^*J*_HH_ = 6.4 Hz, H1′), 5.39 (1H, dt, ^3^*J*_HH_ = 8.8, 5.2 Hz, H3′), 5.30, (1H, t, ^3^*J*_HH_ = 5.2 Hz, OH) 4.28 (1H, dt, ^3^*J*_HH_ = 5.2, 3.6 Hz, H4′), 3.81 (3H, s, O-CH_3_), 3.77-3.64 (2H, m, H5′), 2.83-2.66 (2H, m, H2′), 1.83 (3H, s, CH_3_). ^13^C{^1^H} NMR (100 MHz, DMSO-*d*6) δ 164.2 (C=O), 159.5 (Cq_Ar_), 150.9(C=O), 146.9 (Cq_Ar_), 136.7 (CH_Ar_), 126.9 (2CH_Ar_), 123.6 (CH_Ar_), 120.4 (CH_trz_), 114.8 (2CH_Ar_), 110.1 (CH_Ar_), 84.8 (C4′), 84.3 (C1′), 61.2 (C5′), 59.7 (C3′), 55.6 (s, O-CH_3_), 37.6 (C2′), 12.7 (CH_3_).

### Compound 5

A mixture of 3-azido-3-deoxythymidine (100 mg, 0.375 mmol), 1-octyne (110 μl, 0.748 mmol) and Cp*RuCl(PPh_3_)_2_ (15 mg, 0.05 eq.) were dissolved in dioxane (5 ml) under N_2_ atmosphere. The resulting reaction mixture was stirred at 80°C for 48 hours. After cooling, the solvent was removed under vacuum on rotary evaporator. The solid residue was purified by silica gel chromatography (DCM:MeOH 9:1 v/v). Compound **5** was obtained as a white powder (123 mg). Yield: 87%.^1^H NMR (400 MHz DMSO-*d*6) δ 11.4 (1H, s, NH), 7.80 (1H, s, H_Ar_), 7.57 (1H, s, H_trz_), 6.51 (1H, t, ^3^*J*_HH_ = 6.8 Hz, H1′), 5.34 (1H, s, OH), 5.12 (1H, d, ^3^*J*_HH_ = 4.2 Hz, H3′), 4.22 (1H, d, ^3^*J*_HH_ = 3.5 Hz, H4′), 3.72-3.59 (2H, dd, H5′), 2.70 (2H, t, ^3^*J*_HH_ = 7.6 Hz, H_oct_), 2.62 (2H, t, ^3^*J*_HH_ = 6.3 Hz, H2′), 1.82 (3H, s, CH_3_), 1.61 (2H, m, ^3^*J*_HH_ = 6.5 Hz, H_oct_), 1.30 (6H, m, H_oct_), 0.87 (3H, m, H_oct_). ^13^C{^1^H}NMR (400 MHz, DMSO-*d*6) δ 164.2 (C=O), 150.9 (C=O), 138.3 (Cq_ta_), 136.6 (CH_Ar_), 132.1 (CH_trz_), 110.1 (Cq_Ar_), 85.3 (CH4′), 84.9 (CH1′), 61.6 (CH5′), 57.4 (CH4′), 37.6 (CH2′), 31.3, 28.7, 27.9, 22.5, 22.4 (CH_oct_), 14.4 (CH_oct_), 12.8 (CH_3_).

### Compound 6

A mixture of 3-azido-3-deoxythymidine (100 mg, 0.375 mmol), 2-ethnyl-6 methoxynaphthalene (102 mg, 0.561 mmol) and Cp*RuCl(PPh_3_)_2_ (15 mg, 0.05 eq.) were dissolved in dioxane (4 ml) under N_2_ atmosphere. The resulting mixture was stirred at 60°C for 48 hours. After cooling, the solvent was removed under vacuum on rotary evaporator. The solid residue was purified by silica gel chromatography (DCM: MeOH 9:1 v/v). Compound 3 was obtained as a grey powder (89 mg) Yield: 53%. ^1^H NMR (400 MHz DMSO-*d*_6_) δ 11.4 (1H, s, NH), 8.06 (1H, s, H_Ar_), 7.99-8.01 (2H, m, H_trz_, H_Ar_), 7.93 (1H, d, ^3^*J*_HH_ = 8.96 Hz, H_Ar_), 7.78 (1H, s, H_Ar_), 7.59 (1H, d, ^3^*J*_HH_ = 8.24 Hz, H_Ar_), 7.44 (1H, s, H_Ar_), 7.27 (1H, d, ^3^*J*_HH_ = 8.76 Hz, H_Ar_), 6.60 (1H, t, ^3^*J*_HH_ = 6.9 Hz, H1′), 5.23-5.27 (1H, m, H4′), 4.44 (1H, d, H3′), 3.92 (3H, s, O-Me), 3.49-3.63 (2H, dd, H5′), 2.61-2.67 (2H, dd, H2′), 1.77 (3H, s, CH_3_). ^13^C{^1^H}NMR (400 MHz, DMSO-*d*_6_) δ 164.2 (C=O), 158.7 (Cq_Ar_), 150.9 (C=O), 138.6 (Cq_Ar_), 136.6 (CH_Ar_), 134.9 (Cq_Ar_), 133.4 (CH_trz_), 130.3 (CH_Ar_), 129.0 (CH_Ar_), 128.6 (Cq_Ar_), 128.1 (CH_Ar_), 127.2 (CH_Ar_), 121.7 (Cq), 120.1 (CH_Ar_), 110.2 (Cq), 106.4 (CH_Ar_), 85.4 (CH4′), 85.1 (CH1′), 61.8 (CH5′), 58.8 (CH_3_′), 55.8 (OCH_3_) 38.3 (CH2′), 12.7 (CH_3_).

### Compound 7

A mixture of 3-azido-3-deoxythymidine (150 mg, 0.568 mmol), diphenylacetylene (202 mg, 1.14 mmol) and Cp*RuCl(PPh_3_)_2_ (22 mg, 0.05 eq.) were dissolved in dioxane (3 ml) under N_2_ atmosphere. The resulting mixture was stirred at 60°C for 48 hours. After cooling, Et_2_O was added to reaction mixture and a dark green solid precipitated. Solid was filtered and pentane was then added to the filtrate, precipitating a light green/white solid. This solid was filtered and washed with Et_2_O and pentane before it was let to dry. Compound 5 was obtained as a light green/white powder (146 mg). Yield: 58%. ^1^H NMR (400 MHz DMSO-*d*_6_) δ 11.4 (1H, s, NH) 7.77 (1H, s, H_Ar_), 7.60-7.30 (10H, H_Ar_), 6.59 (1H, t, ^3^*J*_HH_ = 7.8 Hz, H1′), 5.22 (1H, t, ^3^*J*_HH_ = 4.88 Hz, OH), 4.87 (1H, d, ^3^*J*_HH_ = 8.24 Hz, H3′), 4.42 (1H, d, ^3^*J*_HH_ = 1.84 Hz, H4′), 3.58-3.48 (2H, m, H5′), 2.74-2.25 (2H, m, H2′), 1.77 (3H, s, CH_3_). ^13^C{^1^H} NMR (400 MHz, DMSO-*d*_6_) δ 164.1 (C=O), 150.9 (C=O), 143.3 (Cq), 136.5 (1C, CH_3_), 134.6 (Cq), 131.1 (Cq), 130.8–126.7 (11C, CH_Ar_), 127.5 (Cq), 110.2 (Cq), 85.3 (CH4′), 85.2 (CH1′), 61.9 (CH_5′_), 59.1 (CH3′), 37.9 (CH2′), 12.7 (CH_3_).

### Compound 8

Compound **6** (37.4 mg, 0.00832 mmol) was dissolved in DMF (3 ml) under N_2_ atmosphere and methyl iodide (17.6 μl, 0.0283 mmol) was added. The resulting mixture was stirred at 100°C for 24 hours. After cooling to room temperature, Et_2_O was added and an orange oil precipitated. The oily product was continuously washed with Et_2_O until solidification, and then filtered-off and dried under vaccum. Yield: 91%, 59.9 mg. ^1^H NMR (400 MHz DMSO-*d*_6_) δ 11.4 (1H, s, NH), 9.18 (1H, s, H_trz_), 8.19 (1H, s, H_Ar_), 8.12 (1H, d, ^3^*J*_HH_ = 8.40 Hz, H_Ar_), 8.02 (1H, d, ^3^*J*_HH_ = 9.04 Hz, H_Ar_), 7.76 (1H, s, H_Ar_), 7.66 (1H, d, ^3^*J*_HH_ = 8.16 Hz, H_Ar_), 7.52 (1H, s, H_Ar_), 7.34 (1H, d, ^3^*J*_HH_ = 8.68 Hz, H_ar_), 6.47 (1H, t, ^3^*J*_HH_ = 6.76 Hz, H1′), 5.48 (1H, d, ^3^*J*_HH_ = 7.36 Hz, H3′), 5.33 (1H, s, OH), 4.58 (1H, d, H4′), 4.45 (3H, s, N-Me), 3.94 (3H, s, O-Me), 3.60-3.50 (2H, dd, H5′), 2.75-2.50 (2H, dd, H2′), 1.77 (3H, s, CH_3_). ^13^C{^1^H}NMR (400 MHz DMSO-*d*_6_) δ 164.1 (C=O), 159.6 (C_q_), 150.9 (C=O), 143.3 (C_q_), 136.5 (CH_Ar_), 135.9 (C_q_), 130.7 (CH_Ar_), 130.6 (CH_Ar_), 128.6 (CH_Ar_), 128.2 (C_q_), 126.7 (CH_Ar_), 120.7 (CH_Ar_), 117.5 (C_q_), 110.3 (C_q_), 106.6 (CH_Ar_), 85.1 (CH1′), 84.6 (CH4′), 62.5 (CH3′), 61.7 (CH5′), 56.0 (OMe), 40.9 (N-Me), 37.8 (CH2′), 12.7 (CH_3_).

### Compound 9

Compound **4** (42 mg, 0.11 mmol) was dissolved in DMF (2 ml) and iodomethane was added (65 μL, 1.1 mmol). The mixture was stirred at 100 °C for 24 hours. After cooling, Et_2_O was added to precipitate the compound. The solid was isolated by filtration, washed with Et_2_Oand dried under vaccum. Compound **9** was obtained as a brown powder (44 mg, 77.9%). ^1^H NMR (400 MHz, DMSO-d6) δ 11.40 (1H, s, NH), 9.05 (1H, s, H_trz_), 7.77 (1H, d, ^4^J_HH_ = 0.8 Hz, H_Ar_), 7.59 (2H, d, 3*J*_HH_ = 8.8 Hz, H_Ar_), 7.24 (2H, d, ^3^*J*_HH_ = 8.8 Hz, H_Ar_) 6.44 (1H, dd, ^3^*J*_HH_ = 6.4 Hz, H1′), 5.39-5.34 (1H, m, H3′ and OH), 4.515 (1H, dt, ^3^*J*_HH_ = 3.6, 3.2 Hz, H4′), 4.42 (3H, s, N-CH_3_), 3.88 (3H, s, O-CH_3_), 3.64-3.51 (2H, m, H5′), 2.87-2.65 (2H, m, H2′), 1.78 (3H, s, CH_3_). ^13^C{^1^H} NMR (100 MHz, DMSO-*d6*) δ 164.1 (s, C=O), 162.0 (CH_Ar_), 150.8 (s, C=O), 143.0 (Cq_trz_), 136.4 (CH_Ar_), 132.0 (2CH_Ar_), 130.3 (CH_trz_), 115.5 (2CH_Ar_), 114.5 (Cq_Ar_), 110.3 (Cq_Ar_), 85.0 (CH1′), 84.5 (CH4′), 62.3 (CH_3_′), 61.6 (CH5′), 56.0 (O-CH_3_), 40.6 (N-CH_3_), 37.6 (CH2′), 12.7 (CH_3_).

### Peripheral Mononuclear Blood Cells (PBMCs) and infection

PBMCs were isolated from healthy donors by gradient ficoll. Cells were cultured during 7 days in RPMI (Gibco) supplemented with 10% of Fetal Bovine Serum (Biochrom) and 1% of antibiotic-antimicotic (Gibco), in the presence of phytohemagglutinin (0.4 μg/ml) and IL-2 (20 IU/ml). On method 1, cells were infected with GFP-tagged NLENG1-IRES HIV-1 virus, kindly provided by David N. Levy (12), and the AZT derivatives were added to the infected cells at a concentration of 10μM. On method 2, cells were incubated with the AZT derivatives (10 μM) 18 hours before infection with NLENG1-IRES virus. After infection, the AZT derivatives (10 μM) were added again to cells, on both methods. On day 5 post infection, the results were assessed by flow cytometry (BD FACS Canto II). Cells were stained with the viability dye eFluor 780 (eBioscience) and labelled with a PerCP-Cy5.5-conjugated CD4 antibody (Biolegend. All the results were analysed by Flow Jo 10.

## REFERENCES

1 S. Broder, Antiviral Research, 2010, 85, 1–18.

2 P. Zhan, C. Pannecouque, E. De Clercq and X. Liu, J. Med. Chem., 2016, 59, 2849–2878.

3 C. V. Fletcher, K. Staskus, S. W. Wietgrefe, M. Rothenberger, C. Reilly, J. G. Chipman, G. J. Beilman, A. Khoruts, A. Thorkelson, T. E. Schmidt, J. Anderson, K. Perkey, M. Stevenson, A. S. Perelson, D. C. Douek, A. T. Haase and T. W. Schacker, PNAS, 2014, 111, 2307–2312.

4 R. Granich, S. Crowley, M. Vitoria, C. Smyth, J. G. Kahn, R. Bennett, Y.-R. Lo, Y. Souteyrand and B. Williams, Current Opinion in HIV and AIDS, 2010, 5, 298–304.

5 S. Raic-Malic and A. Mescic, Curr. Med. Chem., 2015, 22, 1462–1499.

6 S. K. V. Vernekar, L. Qiu, J. Zhang, J. Kankanala, H. Li, R. J. Geraghty and Z. Wang, J Med Pharm Chem., 2015, 58, 4016–4028.

7 V. R. Sirivolu, S. K. V. Vernekar, T. Ilina, N. S. Myshakina, M. A. Parniak and Z. Wang, J. Med. Chem., 2013, 56, 8765–8780.

8 P. Chittepu, V. R. Sirivolu and F. Seela, Bioorganic & Medicinal Chemistry, 2008, 16, 8427–8439.

9 K. F. Donnelly, A. Petronilho and M. Albrecht, Chem. Commun., 2013, 49, 1145–1159.

10 J. G. Silva, N. P. Martins, R. Henriques and H. Soares, J. Immunol., 2016, 197, 4042–4052.

11 H. Soares, R. Henriques, M. Sachse, L. Ventimiglia, M. A. Alonso, C. Zimmer, M.-I. Thoulouze and A. Alcover, J Exp Med, 2013, 210, 2415–2433.

12 B. Trinité, C. N. Chan, C. S. Lee, S. Mahajan, Y. Luo, M. A. Muesing, J. M. Folkvord, M. Pham, E. Connick and D. N. Levy, PLoS ONE, 2014, 9, e110719-.

